# Wastewater Influent Microbial Immigration and Contribution to Resource Consumption in Activated Sludge Using Taxon-Specific Mass-Flow Immigration Model

**DOI:** 10.1101/2022.08.15.504022

**Authors:** Bing Guo, Chenxiao Liu, Claire Gibson, Nouha Klai, Xuan Lin, Dominic Frigon

## Abstract

Wastewater influent microorganisms are part of the total chemical oxygen demand (COD) and affect the activated sludge (AS) microbial community. Precise modeling of AS processes requires accurate quantification of influent microorganisms, which is missing in many AS models (ASMs). In this study, influent microorganisms in COD unit were determined using a fast quantification method based on DNA yield and was compared with conventional respirometry method. The actively growing influent microorganisms were identified. A mass-flow immigration model was developed to quantify the influent-to-AS immigration efficiency (*m*_*i*_) of specific taxon *i* using mass balance and 16S rRNA gene high-throughput sequencing data. The modelled average *m* was 0.121-0.257 in site 1 (LaPrairie), and 0.050-0.126 in site 2 (Pincourt), which were corrected to 0.111-0.186 and 0.048-0.109 respectively using a constrain of *m*_*i*_ ≤ 1. The model was further developed to calculate contributions to organic substrate consumption by specific taxa. Those genera with zero or negative net growth rates were not completely immigration dependent (*m*_*i*_ < 1) and contributed to 2.4% - 5.4% of the substrate consumption. These results suggest that influent microbiome may be important contributors to AS microbiome assembly and system performance (substrate consumption), which may help to improve future AS process modelling and design.

**Synopsis:** Influent microbial immigration lacks detailed taxon-specific quantification. This study presents quantitative methods and models for influent biomass, mass-flow immigration model, and resource consumption in activated sludge.

**Graphic Abstract:** 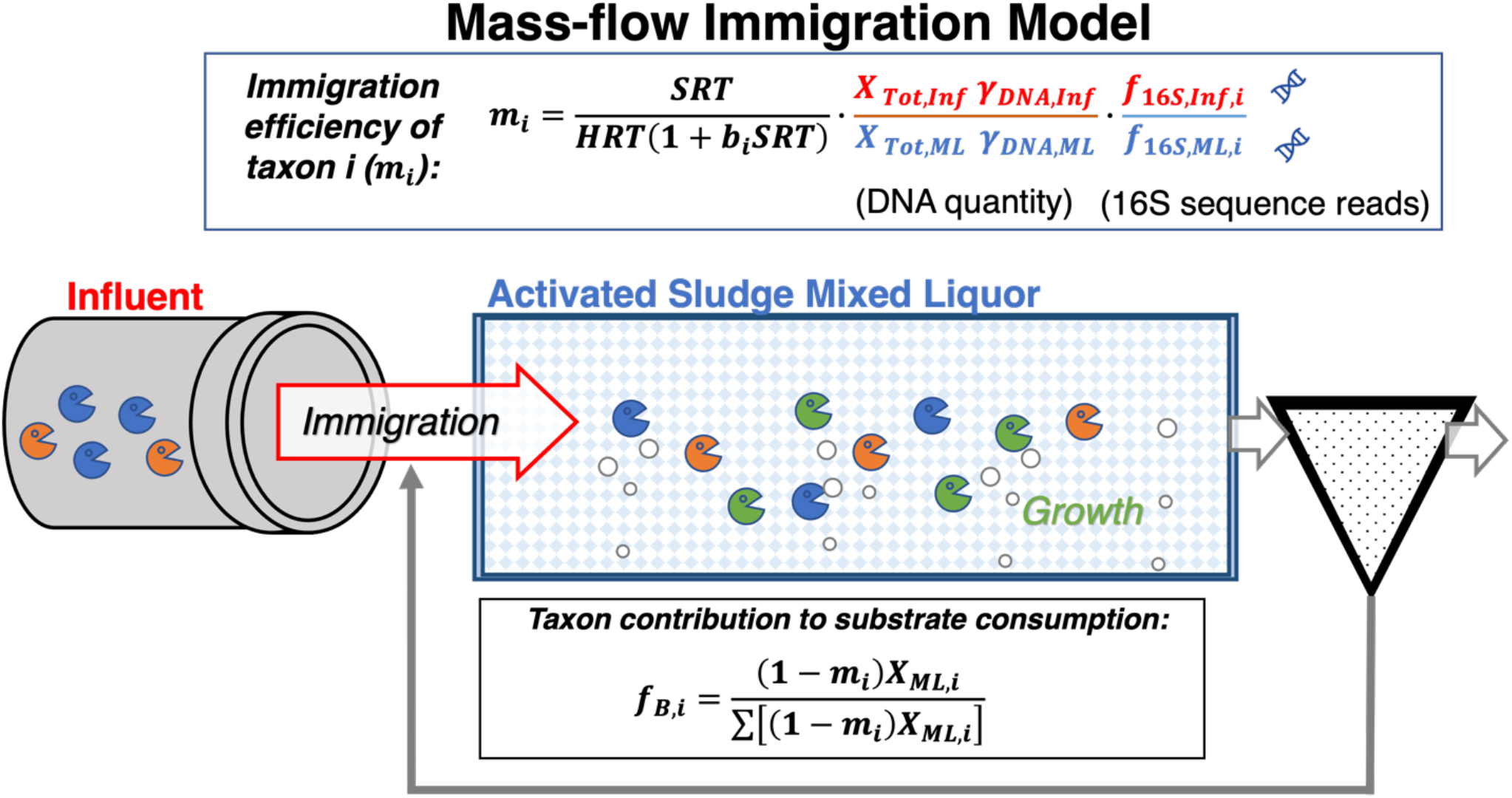

## 1. Introduction

The use of mathematical models to support the design and operation of water resource recovery facilities (WRRFs) has matured to efficiently tackle problems of process optimization with respect to energy consumption and operation costs, development of control strategies, and improvement of effluent quality.^1^ However, effective utilization of these models requires detailed calibration and validation, which remains a difficult task to accomplish, given the lack of detailed information on influent characteristics and the interdependency between model parameters and influent description.^1-3^ Influent biomass is an emerging key influent chemical oxygen demand (COD) fraction,^4^ which is gaining importance with the interest in activated sludge systems operated at very high rates (i.e. solids residence time [SRT] < 3 days)^5, 6^ or at low temperatures with microbial activities maintained near or below washout SRTs such as nitrification activity during winter.^7^

The effects of influent microbiome input to the activated sludge processes have drawn interests on general heterotrophic organisms ^8-10^ and specific functional microbes such as nitrifiers and polyphosphate accumulating organisms (PAO).^7, 11-13^ Until now, detailed influent microbiome input has largely been neglected in conventional process modeling frameworks, e.g., activated sludge models (ASMs). It has been documented that the nitrifying bacterial populations have similar structure in the influent and the activated sludge mixed liquor.^7^ Further, the input of nitrifiers from the influent has been shown to effectively maintain and restore activated sludge nitrification at low temperatures,^7^ and to shorten the necessary SRT in cold regions.^14^

In the case of heterotrophic population, researchers and engineers have been interested in more detailed modeling approaches capable to account the impact of influent heterotrophs on unit process performance.^3, 15^ Part of this need arose from modeling exercises of high-rate (low SRT) A-stage wastewater treatment processes, where influent heterotrophs can contribute to significant soluble COD removal.^16^ However, the fraction of influent COD attributed to active heterotrophs is widely ranging from 2 to 30%.^17^ Therefore, there is a need to develop appropriate protocols to evaluate the quantities of influent heterotrophs which should be considered affecting activated sludge systems.^4^

Using 16S rRNA gene amplicon sequencing, the resolution of microbial community composition has been greatly improved. Meanwhile, one of the major concerns with the current modeling approach of lumping all heterotrophs in a single population is the lack of consideration for differences among species, namely the selection effect and the immigration (or transfer) efficiency between the influent and the activated sludge microbial communities.^18^ This issue needs to be resolved if one wants to use microbial community composition data to inform the modeling exercise. A few studies used gene sequencing to evaluate the neutrality vs. the selection of heterotrophic populations during the immigration from influent to activated sludge. Using species rank and mass balance, the efficiency of microbial immigration was assessed, suggesting that some populations were highly selected by niche partitioning while others were well maintained in the activated sludge community by immigration from influent.^19-23^ The specific SRT appears to play a significant role in defining the efficiency of immigration. Indeed, in high-rate activated sludge processes, microbial communities showed more neutral assembly (i.e., lower level of selections between the influent and the mixed liquor) than in low-rate processes, implying that immigration has stronger influences on the microbial composition at low SRTs.^5^ Despite these key observations, the methods used in these studies to characterize the microbial communities, unfortunately, do not explicitly quantify the variation in growth activity exhibited by each heterotroph, and the data are not in units and formats usable by activated sludge modelers.

To understand the impact of influent microbiome to activated sludge microbiome and system performance in a quantifiable way, it is essential to generate a modelling tool that utilises microbial community data and operational parameters. This study aims at: (1) Quantifying the fraction of active microbiome in influent and the effects on activated sludge microbiome; (2) Modeling the net growth rates and immigration efficiencies of microorganisms using common units (COD); (3) Evaluating the contribution to resource consumption by high immigration and low growth genera. Conventional respirometry test and high throughput 16S rRNA gene amplicon sequencing were used for total and detailed heterotrophic biomass estimation.

## 2. Materials and methods

### 2.1. Influent sampling and respirometry test

Influent and activated sludge mixed liquor samples were taken from LaPrairie and Pincourt WRRFs (Québec, Canada). Plant configuration and operational parameters are shown in Table S1. The average temperature of activated sludge was 11 °C. Influent solids were settled from approximate 30 L of wastewater, decanted, then resuspended in the supernatant to reach 9×, 6×, 4× and 1× of original solids concentration to provide enough sensitivity for the respirometry tests. Respirometry tests were performed with a model AER-200 respirometer (Challenge Technology, USA) at ambient temperature (24 °C). Rate estimation was corrected for 20 °C and 11 oC using *r* = *r*_20°C_ × 1.029^(*T*−20°C)^ and *r* = *r*_11°C_ × 1.029^(*T*−11°C)^.

### 2.2. Calculation of activate influent heterotrophic biomass fraction

Estimation of active ordinary heterotrophic organisms (OHO, notations from ASMs^24^) concentration (*X*_*OHO*_) in influent were calculated (Eq. 1) using the linear regression of the natural log of oxygen uptake rate (OUR) over time during the exponential growth^25^ (Figure S1, insert).

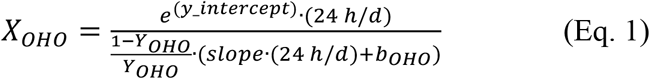

where *X*_*OHO*_ is the active heterotrophic organisms in influent, *Y*_*OHO*_ is the default growth yield (0.666 mg-COD_OHO_/mg-COD_substrate_), *b*_*OHO*_ is the default decay rate.

A second method to determine the active heterotrophic biomass fraction in the influent (*f*_*OHO,Inf*_) is using DNA yield (*γ*_*DNA*_, Eq. 2). The DNA yield is measured by extraction and quantification using absorption at 260 nm. To convert the DNA units into COD units, it is assumed that the influent and activated sludge mixed liquor (ML) biomass have same DNA/cell ratio.

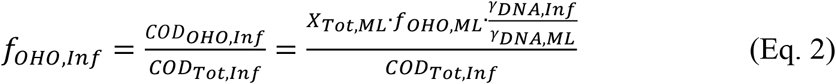

where *f*_*OHO,Inf*_ and *f*_*OHO,ML*_ are heterotrophic fractions in the influent and mixed liquor, respectively; *COD*_*OHO,Inf*_ and *COD*_*Tot,Inf*_ are the influent heterotrophic biomass COD and total COD concentrations (mg-COD/L); *γ*_*DNA,Inf*_ and *γ*_*DNA,ML*_ are DNA extraction yield from influent and mixed liquor biomass (μg-DNA/mg-solid); *X*_*Tot,ML*_ is the mixed liquor total solids concentration (mg-COD/L).

The mixed liquor active biomass fraction and decay rate were estimated using respirometry test ^26^(supplementary information).

### 2.3. 16S rRNA gene amplicon sequencing and bioinformatics analysis

During respirometry test, biomass samples were harvested at the end of the exponential growth phase, centrifuged, and stored at −80 °C before DNA extraction. Genomic DNA was extracted using DNeasy PowerSoil Kit (Qiagen Inc., Toronto, Canada) following the manufacturer’s instructions. PCR was performed using modified 515F and 806R primers^27, 28^ targeting the V4 region of 16S rRNA genes. Details of PCR materials and conditions are shown in Table S2. Amplicons were sequenced on Illumina MiSeq PE250 platform at McGill University and Génome Québec Innovation Centre (Montréal, QC, Canada). The raw sequences were quality-filtered using DADA2^29^ in Qiime2 pipelines.^30^ Taxonomy was assigned using GreenGenes (version gg_13_8) reference database at 99% similarity^31, 32^ and tabulated at genus level. For the immigration analysis, genus-level taxa were used to minimize signals caused by within-genus neutrality.^33^ Microbial community alpha and beta diversities, Bray-Curtis distance and Principal Coordinate Analysis (PCoA) were analyzed using R “vegan” package.^34^

### 2.4. Mass-flow immigration model

We report here the key equations used to calculate the overall and taxon-specific mass-flow immigration efficiency. The following equations were derived to quantify the amount of mixed liquor heterotrophic biomass (*X*_*OHO,ML*_) immigrated from the influent (*X*_*OHO,Inf*_) and the amount produced from growth (the consumption of substrate resources), *Y*_*OHO*_(*XC*_*B*_ + *S*_*B*_)_*Inf*_, where *Y*_*OHO*_ is the heterotrophic growth yield, *XC*_*B*_ is the slowly biodegradable substrates, *S*_*B*_ is the soluble biodegradable substrates and residual substrate is assumed negligible.

From steady-state mass balances on biomass and on substrates, *X*_*OHO,ML*_ can be found using Eq. 3:^26^

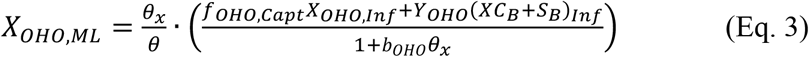

where *θ*_*x*_ is solids retention time (SRT); *θ* is hydraulic retention time (HRT); *f*_*OHO,Capt*_ is the fraction of influent biomass captured by the mixed liquor flocs; *X*_*OHO,Inf*_ and *X*_*OHO,ML*_ are the active heterotrophic biomass in influent and mixed liquor; *Y*_*OHO*_ is the heterotrophic growth yield; *b*_*OHO*_ is the heterotrophic decay rate.

Mass-flow immigration efficiency (*m*) is defined as the proportion of biomass in the mixed liquor derived from the influent immigration to the total biomass^18^ and expressed by Eq. 4:

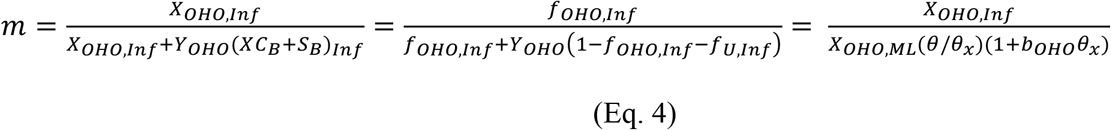

where *m* is the mass-flow immigration efficiency; *XC*_*B*_ is the slowly biodegradable substrates; *S*_*B*_ is the soluble biodegradable substrates; *f*_*U,Inf*_ is the undegradable fraction of influent COD.

Using 16S rRNA gene amplicon sequencing results, the proportion of specific taxon *i* in the biomass sample (*f*_16*S,i*_) can be quantified. The biomass concentration of the taxon *i* can be defined by Eq. 5:

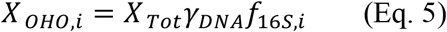

where *X* _*OHO,i*_ is the biomass concentration of the taxon *i*; *X* _*Tot*_ is the total concentration of solids; *γ*_*DNA*_ is the mass of DNA per solids; *f*_*DNA,i*_ is the proportion of the taxon *i* in total 16S rRNA gene amplicon sequence reads.

Consequently, from Eq. 3, 4 and 5, mass-flow immigration of the taxon *i* (*m*_*i*_) is defined by Eq. 6:

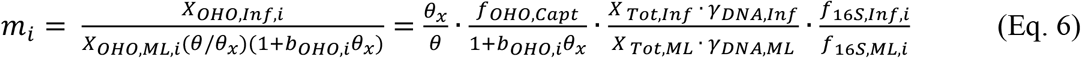

where *m* _*i*_ is the mass-flow immigration efficiency of the taxon *i*; *X* _*OHO,Inf,i*_ and *X* _*OHO,ML,i*_ are the biomass concentrations of the taxon *i* in influent and mixed liquor; *b*_*OHO,i*_ is the decay rate; *X* _*Tot,Inf*_ and *X* _*Tot,ML*_ are the total concentrations of solids in influent and mixed liquor; *γ*_*DNA,Inf*_ and *γ*_*DNA,ML*_ are the masses of DNA per solids in influent and mixed liquor; *f*_*DNA,Inf,i*_ and *f*_*DNA,ML,i*_ are the proportions of the taxon *i* in total 16S rRNA gene amplicon sequence reads in influent and mixed liquor.

### 2.5. Resource consumption

Rearranging Eq. 6, a relationship could be identified between the proportions of the taxon *i* in the mixed liquor and in the influent as determined by 16S rRNA gene amplicon sequencing (Eq. 7)

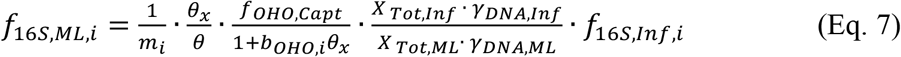

Log transformation results to:

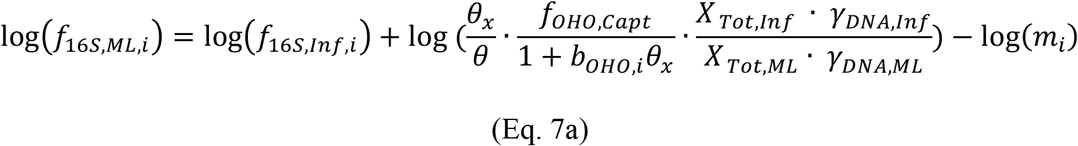

In the case of *m*_*i*_ = 1:

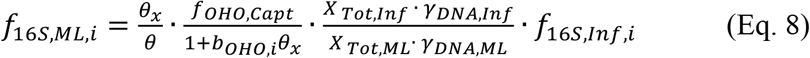

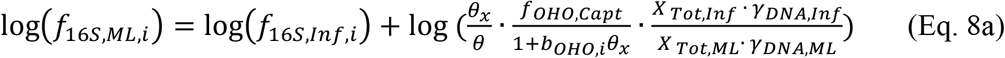

In the case of zero net growth rate *μ*_*OHO,Net,i*_ = 0:

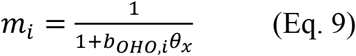

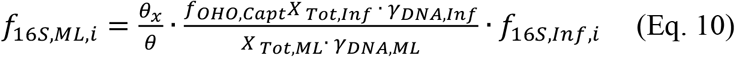

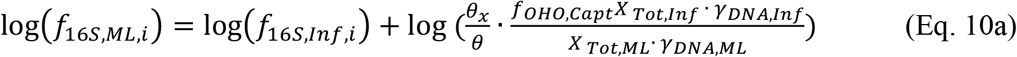

Eq. 8a and Eq. 10a can be visualized on the biplot of log(*f*_16*S,ML,i*_) *vs* log(*f*_16*S,Inf,i*_). The taxa that fall between the two lines of *μ*_*OHO,Net,i*_ = 0 and *m*_*i*_ = 1 indicate that they are not completely relying on immigration (*m*_*i*_ < 1) and still grow and consume substrates in activated sludge.

Finally, the model can be developed to estimate the proportion of heterotrophic resources consumed by the taxon *i*. This is important because a negative growth rate does not mean zero consumption of any resources.

Derived from Eq. 3 assuming *f*_*OHO,Capt*_ = 1:

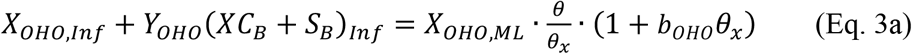

Derived from Eq. 4:

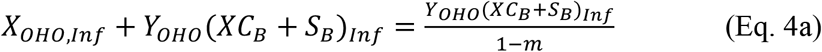

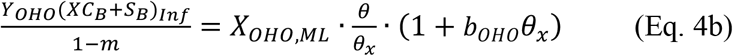

Thus, the amount of substrate resources consumed by the taxon *i*, is estimated in Eq. 11:

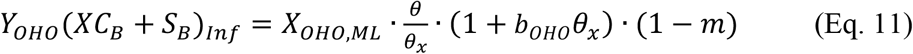

Similarly, for taxon *i:*

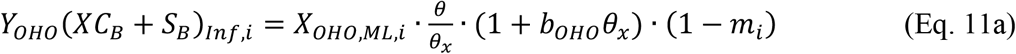

Assuming similar yields and decay rates, the proportion of substrates consumed by the taxon *I* (*f*_*B,i*_) is calculated by Eq. 10:

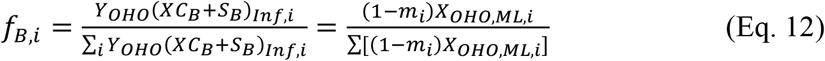

### 2.6. Statistical analysis

To test the growth of influent microorganisms in the respirometry experiment, the rank relative abundances of a taxa and the ranks (1, 2, 3 and 4) of food-to-organism (F/M) ratio at different solids concentrations (9×, 6×, 4×, and 1×) were tested using linear regression (Spearman’s correlation) in R, and significance level was indicated at p<0.05. The PCoA Eigen values and dilution factor of F/M (1/9, 1/6, 1/4, and 1) were tested using linear regression in R. Standard deviation propagation is explained in Supplementary Material. The 95% confidence intervals were calculated using propagated standard deviation.

## 3. Results and discussion

### 3.1. Measuring influent heterotrophic biomass fraction

The fraction of active heterotrophic biomass in influent (*f*_*OHO,Inf*_) was assessed using both the respirometry test (Eq. 1)^25^ and by the DNA yield method (Eq. 2). The low sensitivity of the respirometer required the influent solids to be concentrated to 6× or 9× to be able to measure the exponential growth phase in the oxygen uptake rate (OUR) curve (Figure S1). Using this conventional method (Eq. S1), the evaluation of *f*_*OHO,Inf*_ by respirometry relies on the heterotrophic decay rate (*b*_*OHO*_). Depending on the activated sludge model (ASM) used, the value of *b*_*OHO*_ changes dramatically. ASM1 (*b*_*OHO*_=0.62 d^-1^)^35^ uses the lysis-regrowth description and ASM3 (*b*_*OHO*_=0.2 d^-1^) uses the linear maintenance description.^24^ Although the two models lead to the same results with respect to mass balance modelling, there is no guarantee that the individuals of a given species will consume the substrate COD resources freed by the lysis of the same species. Consequently, the choice of decay mechanisms has a profound implication on the values of biomass and immigration efficiencies reported. For this study, we used both values from ASM1 and AMS3 in the calculations allowing comparison.

The calculated influent heterotrophic biomass fraction (*f*_*OHO,Inf*_) from respirometry test and from DNA yield measurements are shown in Table 1. In the site of LaPrairie, the two sampling dates showed large variations of influent COD (172 and 272 mg/L) and total suspended solids (118 and 226 mg/L), and the influent heterotrophic biomass fraction also showed higher values in the latter date (4.85% and 7.34%) using DNA quantification method. The respirometry test method resulted similar trend (higher values in sampling date 2 than date 1), but the estimated *f*_*OHO,Inf*_ values were lower than those by DNA quantification in date 1 and higher in date 2. The impact of decay rate (*b*_*OHO*_) used on the estimation of *f*_*OHO,Inf*_ by respirometry test is clearly seen, where lower *b*_*OHO*_ values in ASM3 resulted in higher biomass fraction values. In the site of Pincourt, influent characteristics also varied but not as much as LaPrairie. The first sampling date showed higher COD, TSS, and *f*_*OHO,Inf*_ by all methods than the second sampling date. The *f*_*OHO,Inf*_ estimated by DNA quantification was 4.25% and 2.06%, both lower than LaPrairie. The respirometry test resulted higher *f*_*OHO,Inf*_ values than the DNA quantification method for both dates and both decay rates in ASM1 and ASM3. Certain variability was shown among sampling site and days while the study did not aim to test the temporal or site differences as shown in other studies^8, 19^ but rather to validate the methods for testing *f*_*OHO,Inf*_. Nonetheless, the measured *f*_*OHO,Inf*_ are within comparable ranges to values reported in the literature.^17^

**Table 1.**
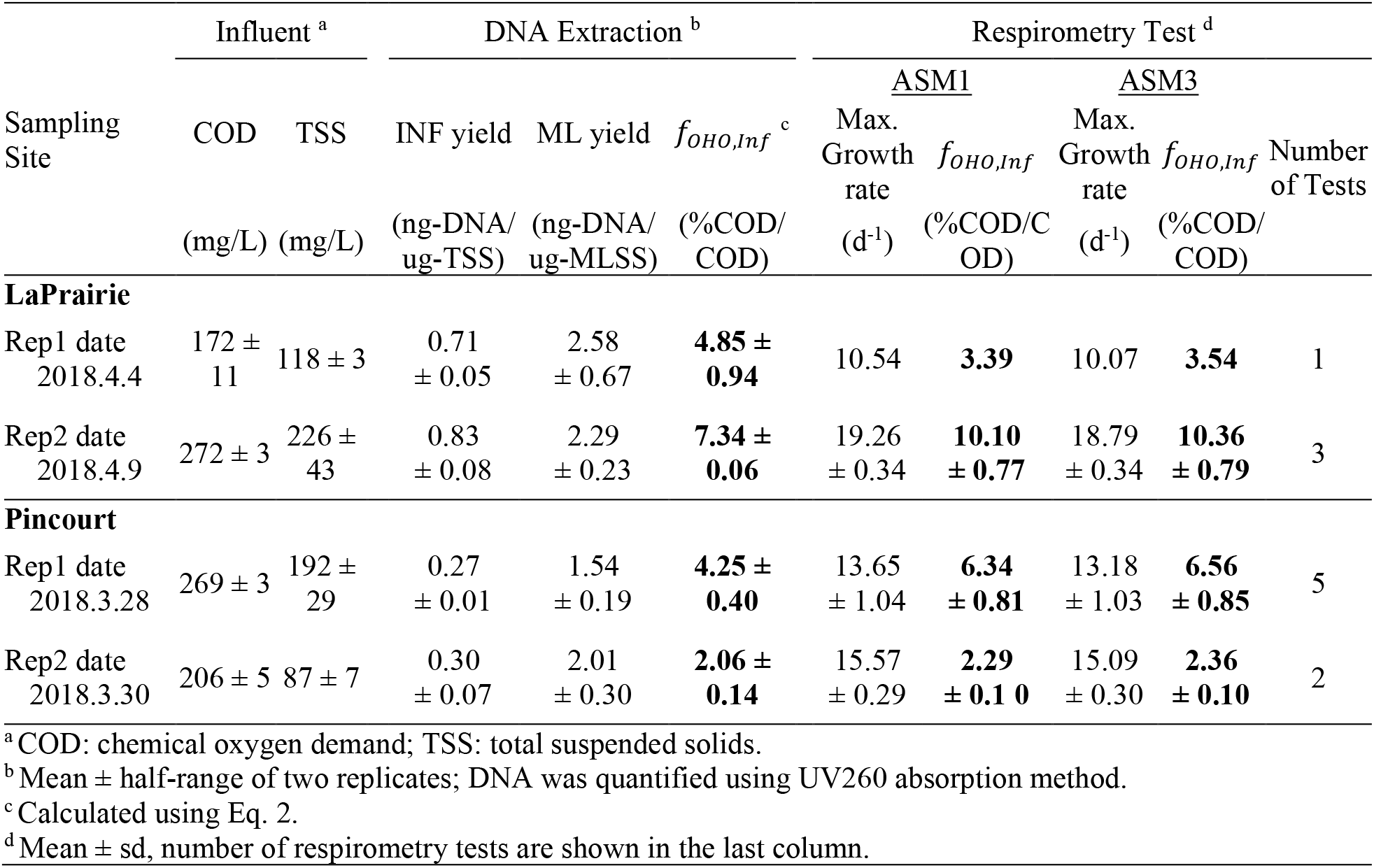
Influent sample characteristics and biomass fractions estimated using DNA extraction yield and using respirometry test.

### 3.2. Microorganisms grown in influent respirometry test

During the respirometry test on influent biomass, microbial growth was observed based on the oxygen update rate. Figure 1a shows that at phylum level, *Proteobacteria, Bacteroidetes* and *Firmicutes* were the most abundant taxa in influent microbiome. An increase in *Proteobacteria* relative abundance was observed after OUR test compared to raw influent, the increase was also observed along the solids concentrating factor from 9× to 1×. With lower influent solids concentration, the food to organism ratio is higher, potentially selecting the fast-growing bacteria, i.e., *Proteobacteria* at phylum level. *Bacteroidetes* slightly decreased with higher F/M ratio for both LaPrairie and Pincout. *Firmicutes* increased with higher F/M ratio in LaPrairie sampling date 1, but decreased in date 2, and increased in Pincourt both dates.

**Figure 1.**
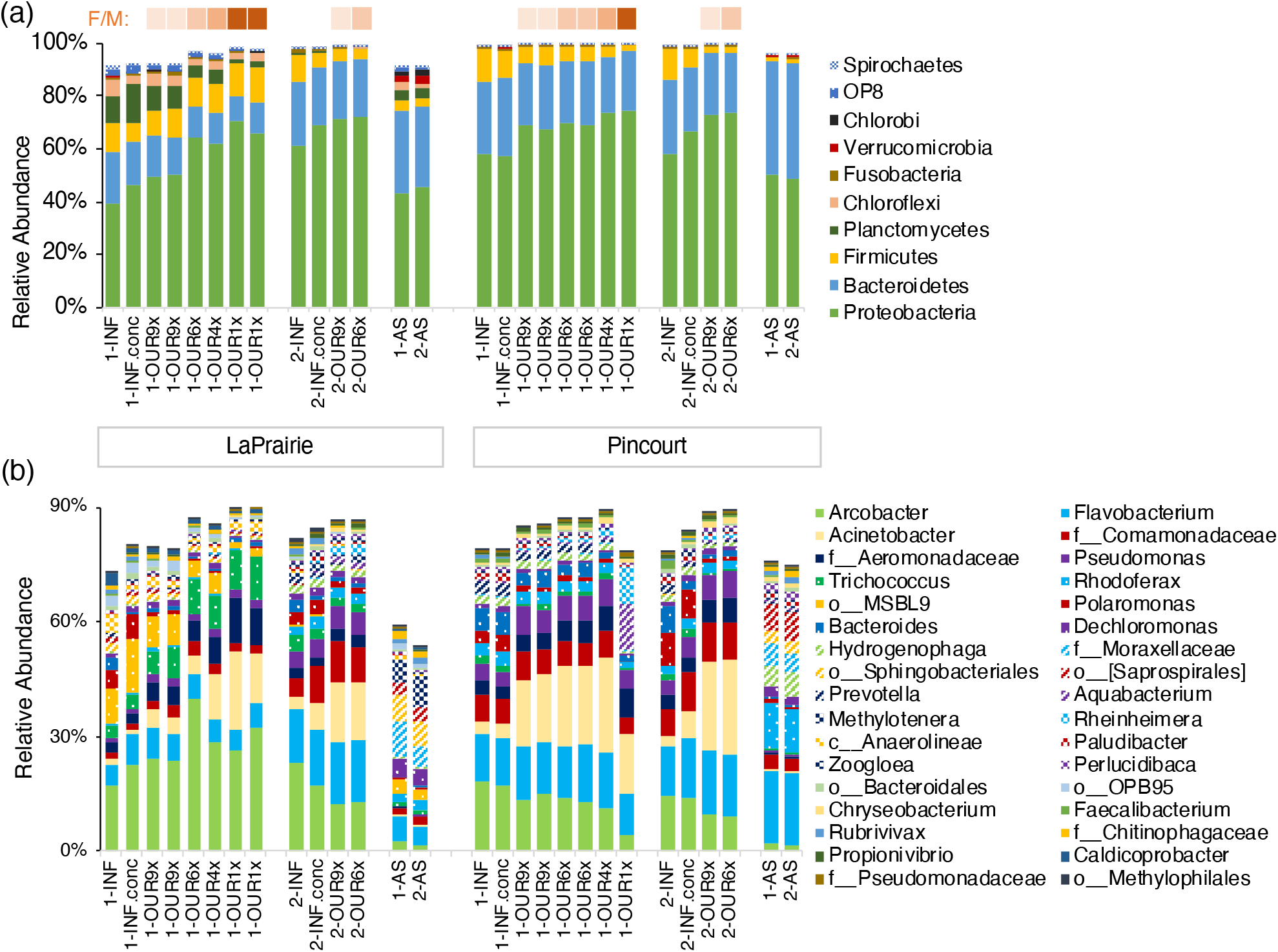
Relative abundance of taxa. (a) top 10 phyla and (b) 50 most abundant genera in influent solids (INF), concentrated influent solids (INF.conc), influent solids grown in respirometry test (OUR) at solids concentrating factors of 9×, 6×, 4×, and 1× in two sampling dates (1 and 2), and activated sludge (AS). F/M ratio is reversely correlated with solids concentrating factor and indicated by color code.

At genus level, the most abundant genera include *Arcobacter, Flavobacterium, Acinetobacter* and *Pseudomonas* (Figure 1b), accounted for 35-74 % of total abundance of influent community and 7-22 % of activated sludge community. Different abundant genera were shown in activated sludge. An uncultured genus in the family *Moraxellaceae* was highly abundant, taking 5-10 % in DNA. *Methylotenera* was especially abundant in LaPrairie, taking 6-9 % of activated sludge. *Methylotenera* has been recognized as an active member in activated sludge at LaPrairie,^36, 37^ even though they were suggested to be slow growers in natural environments.^38^

*Acinetobacter* showed clear increasing trend with higher F/M ratio for both sites, while many genera showed varied abundances. To identify genera that increased in relative abundance during respirometry tests, Spearman’s correlation was performed for taxa at genus level using relative abundances and rank of F/M ratio (increase with decreasing concentration factors 9×, 6×, 4×, and 1×, coded into dummy numbers 1, 2, 3 and 4, respectively). An example with *Acinetobacter* is shown in Table S3 and Figure S2. The LaPrairie and Pincourt regression models resulted in variated coefficient values. Genera with positive coefficients (indicative of active growth) were pooled together, and shown for the most abundant 30 genera (Figure 2). Genera failed for identification (unclassified) were not shown.

**Figure 2.**
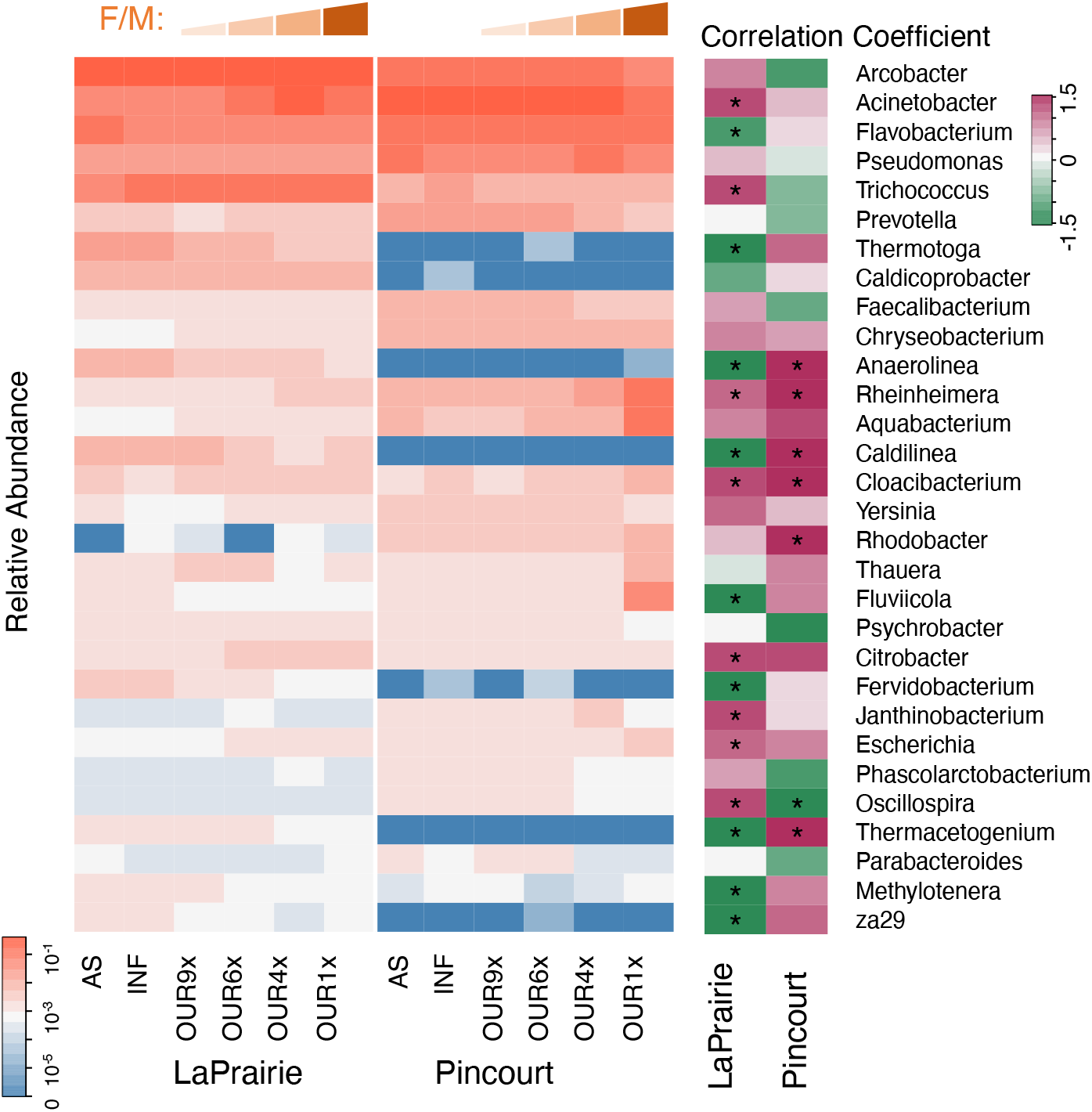
Genera grown in respirometry test (OUR) in either LaPrairie or Pincourt. Spearman’s correlation was performed using relative abundance of genera and rank (1, 2, 3, 4) representing increasing F/M ratio (decreasing solids concentrating factors 9×, 6×, 4×, 1×, respectively). Significance level labelled with * indicates <0.05. Genera from the growth pool data with highest 30 average relative abundance in influents are shown.

Different trends were observed between the two sampling sites’ influent microbial communities. Positive coefficients indicated growth during the respirometry test, which were found for *Rheinheimera* and *Cloacibacterium* in both sites. *Arcobacter, Trichococcus*, and *Acinetobacter* showed positive regression coefficients (growth) in LaPrairie influent respirometry test, but negative in Pincourt (decay). *Fluviicola, Aquabacterium*, and *Cellvibrio* showed an increase in Pincourt influent, but a decrease in LaPrairie. The growth dynamics varied between the sites, possibly due to different wastewater compositions, and variation in microbial activity status.

Some of the influent abundant genera have been considered as decaying bacteria in activated sludge, e.g. *Arcobacter, Bacteroides, Pseudomonas* and *Acinetobacter*.^20^ However, in the current respirometry tests, some of them showed positive growth rates, especially *Acinetobacter* in LaPrairie influent, indicating potential growth in activated sludge. These detailed growth dynamics of influent heterotrophs could be incorporated in precise modeling of activated sludge systems in future studies.

### 3.3. Microbial community diversity

Influent microbial growth during the respirometry test reduced the alpha diversity richness (number of genera) at the end of the test in the lowest F/M ratio in both LaPrairie and Pincourt (Figure S3). When F/M ratio increased, the richness increased in both sites. Due to heterogenous growth among different taxa, it may have caused reduction or elimination of some rare microorganisms with lower F/M due to competition, resulting in reduction in alpha-diversity.

To compare with activated sludge microbial community, the shared genera number and relative abundance are shown in Figure 3. Between LaPrairie influent and activated sludge microbial communities, 336 genera were shared and 210 genera were shared in Pincourt (Figure 3a-b), which took up 71% and 82% of the activated sludge total number of genera, and 72% and 51% of the influent total number of genera in LaPrairie and Pincourt respectively. The shared genera abundance accounted for 94% (LaPrairie) and 98% (Pincourt) of the activated sludge 16S rRNA gene sequence reads (Figure 3c). Strong links between influent and activated sludge communities were implied by the high number and abundance of shared genera.

**Figure 3.**
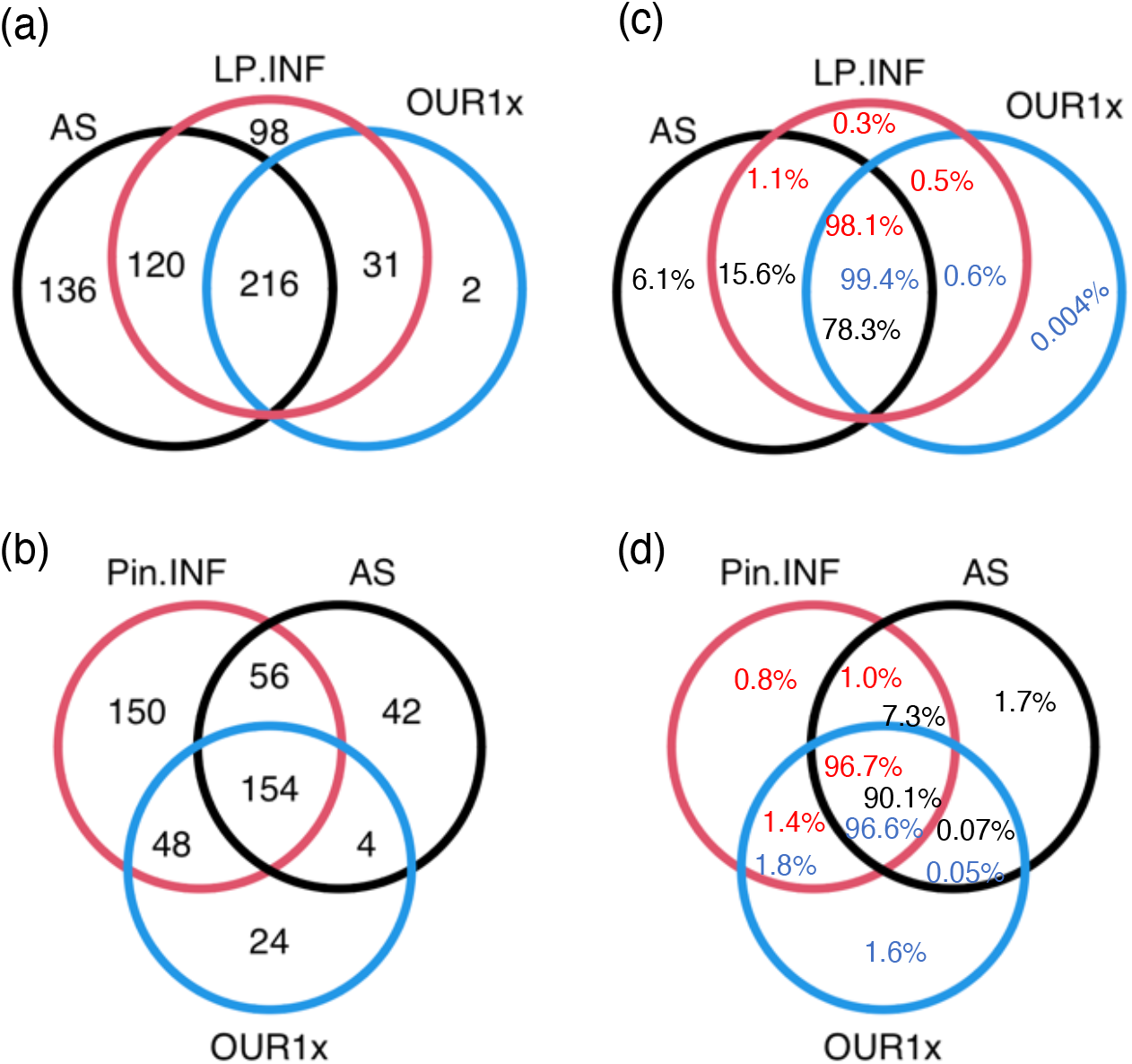
Shared genera in influent (INF), influent biomass after respirometry test (OUR1×), and activated sludge (AS). (a) number of shared genera in LaPrairie (LP); (b) number of shared genera Pincourt (Pin); (c) relative abundances of shared genera in LaPrairie (LP); (d) relative abundances of shared genera in Pincourt (Pin). Colored texts indicate the relative abundance of taxa in influent (red), OUR×1 (blue) and AS (black).

After incubation in respirometry test, only part of genera in raw influent were detected in the highest F/M ratio test (OUR 1×), 247 genera shared and 218 not shared in LaPrairie influent, and 202 shared and 216 not shared in Pincourt influent. The shared number of genera between activated sludge and after growth influent (OUR 1×) reduced to 216 and 206, and 78% and 91% of activated sludge abundance in LaPrairie and Pincourt respectively. Interestingly, 120 genera in LaPrairie and 56 genera in Pincourt were shared only between activated sludge and raw influent, meaning that they disappeared in respirometry test from influent and re-appeared in activated sludge. These genera may have immigrated from influent to activated sludge and were detected in 16S rRNA gene sequencing, but they may not be the active genera because they disappeared (not detectable) in the growth condition even with highest F/M ratio. These results highlight the importance of investigating the fate of specific taxa from influent to activated sludge.

Microbial community beta-diversity of the influent and activated sludge are shown on Principal Coordinate Analysis (PCoA), revealing clear distances along the PCoA1 axis (Figure 4). Growth in the respirometry test drove the community shifting upward along the PCoA2 axis. The communities grown in the highest F/M ratio located the furthest from the original influent communities. Influent biomass after growth shifted towards opposite direction of activated sludge communities. This shift indicated that microbial populations grown in the batch test were dissimilar with the microbial community in full-scale aeration basin. These data clearly demonstrated the differences in community structures between the original influent biomass, the influent biomass after growth and the activated sludge. Consequently, the direct evaluation of the general immigration effects using the total heterotrophic biomass in the influent is inaccurate, the specific community members behave differently in terms of growth and immigration.

**Figure 4.**
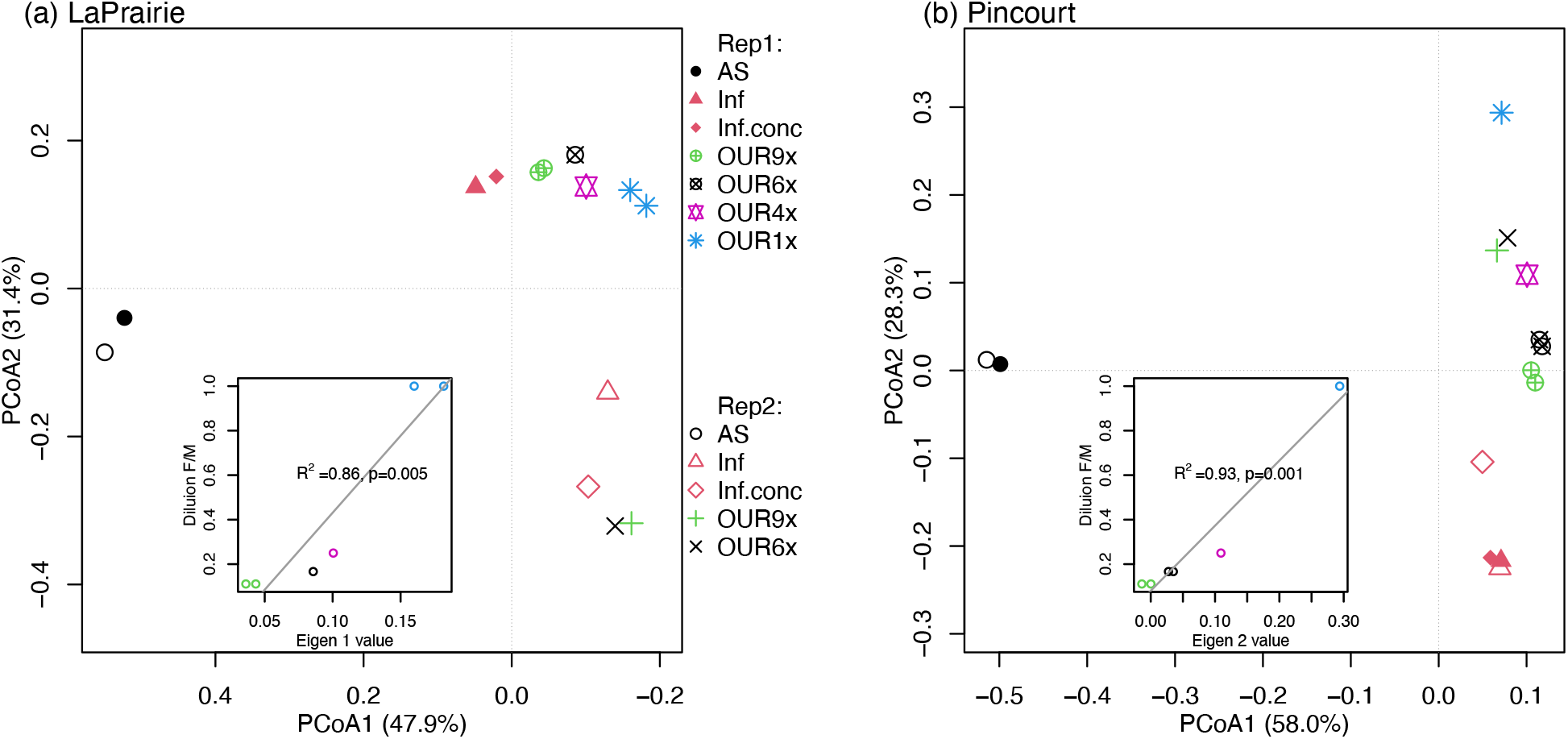
Principal Coordinate Analysis using Bray-Curtis distance of activated sludge (AS), influent (Inf), concentrated influent solids (Inf.conc), influent solids after respirometry tests initiated at different solids concentration factors (OUR9×, 6×, 4×, 1×). (a) LaPrairie, insert graph shows linear regression of Eigen 1 values and microbial community Rep1 after growth (OUR9×, 6×, 4×, 1×); (b) Pincourt, insert graph shows linear regression of Eigen 2 values and microbial community Rep1 after growth (OUR9×, 6×, 4×, 1×).

### 3.4. Mass-flow immigration efficiency of specific taxa

The mass-flow immigration efficiency of specific taxa can be calculated using Eq. 6. The distribution of immigration efficiencies shows that some of the taxa (genus level) had high immigration efficiencies equal to or higher than 1 (Figure S4), indicating that they are not growing in the activated sludge and rely completely on immigration from influent. These genera can be visualized on the log-log activated sludge-influent relative abundance plot under the ***m***_***i***_ = **1** line (Eq. 8a). Figure 5 shows the *m*_*i*_ = 1 reference lines of LaPrairie and Pincourt using the average of two sampling dates. A number of genera were below the maximum immigration *m*_*i*_ = 1 reference lines, and may exhibit either excess decay (i.e., larger *b*_*OHO*_) or low capture rater (*f*_*OHO,Capt*_ < 1). The zero net growth rate reference lines (*μ*_*OHO,Net,i*_ = 0) are also shown in Figure 5 c-d, those genera below the lines indicate zero and negative growth. It should be noted that between the two lines of *μ*_*OHO,Net,i*_ = 0 and *m*_*i*_ = 1, these genera rely partially on immigration (*m*_*i*_ < 1) and slow growth and consume substrates in activated sludge (section 3.5).

**Figure 5.**
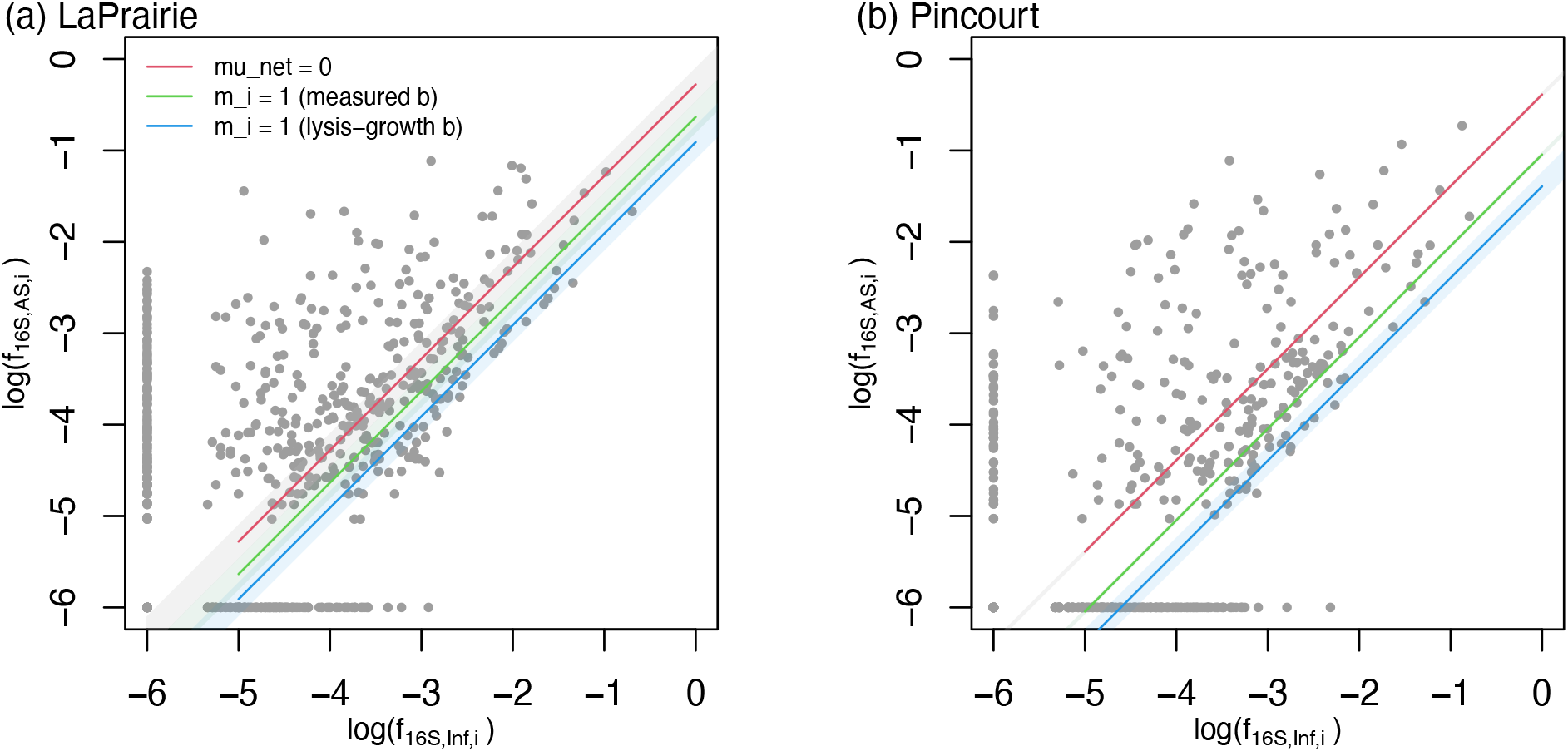
Mass-flow immigration efficiencies of specific taxa at genus level plotted on genus abundance in the activated sludge vs. influent (a) LaPrairie AS vs. Influent; (b) Pincourt AS vs. Influent. Solid lines represent zero net growth rate (Eq. 10a), dash lines represent immigration efficiency of 1 (Eq. 8a). The points at *x* = −6 represent non-detected genera in influent samples; the points at *y* = −6 represent non-detected genera in AS samples. Shaded areas indicate 95% of confidence intervals.

Genera with a relative abundance higher than 1% before or after the respirometry tests were usually detected in activated sludge. However, the relative abundances of genera varied greatly between the influent and activated sludge communities, indicating a strong selection effect. The heterotrophic biomass fractions in the influent (*f*_*OHO,Inf*_) determined using respirometry and DNA quantification methods can be validly used for modeling if the compositions of the influent and mixed liquor communities are the same. Otherwise, the biomass immigration has to be considered differently for different taxa and the *f*_*OHO,Inf*_ should be corrected to represent the selection effect of heterotrophic taxa in the activated sludge.

Based on the distribution of genus-specific immigration efficiencies, the average immigration efficiency was calculated and compared to values assuming 100% efficient immigration efficiency (i.e., neutral or without selection) determined using Eq. 4. The immigration efficiency of several genera were beyond 1 (i.e., genera below the *m*_*i*_ = 1 lines in Figure 5). The immigration efficiencies of these genera were corrected to be 1, and the average immigration efficiencies should be lowered by this correction (Table 2).

**Table 2.** Average immigration efficiency of total microbial community at different decay rates.

| Decay rate | LaPrairie |  |  |  | Pincourt |  |  |  |
| --- | --- | --- | --- | --- | --- | --- | --- | --- |
| | Average Immigration efficiency | | $f_{OHO,Inf}$ (%COD/COD) | | Average Immigration efficiency | | $f_{OHO,Inf}$ (%COD/COD) | |
| | From DNA <sup>a</sup> | Corrected by $m^b$ | From DNA <sup>c</sup> | Corrected by $m^b$ | From DNA <sup>a</sup> | Corrected by $m^b$ | From DNA <sup>c</sup> | Corrected by $m^b$ |
| Measured Decay | 0.232 | 0.173 | 6.095 | 4.545 | 0.090 | 0.083 | 3.155 | 2.910 |
| Lysis-growth decay | 0.123 | 0.113 |  | 5.593 | 0.040 | 0.039 |  | 3.076 |

<sup>a</sup> Calculated from Eq. 4.<sup>b</sup> Cap $m_i$ at maximum value of 1, calculated from Eq. 6.<sup>c</sup> Calculated from Eq. 2.

The decay rate (*b*_*OHO*_) values had a major impact on the calculated immigration efficiencies, with a higher decay rate resulting in a lower immigration efficiency (Table 2). Given that the average immigration efficiency based on the correction of *m* is lower that what could be directly calculated from either the DNA mass or the respirometry test results (Table 2), the biomass fraction in the influent (*f*_*OHO,Inf*_) should be corrected by lowering its value for accurate model simulations of lumped heterotrophs in ASMs. Depending on the systems and the decay rate assumed, the corrected *f*_*OHO,Inf*_ was below the value determined by DNA. The correct *f*_*OHO,Inf*_ declined by 1.55% of total influent COD using the measured decay in ASM3 and by 0.49% using the lysis-growth decay in ASM1 for LaPrairie, and 0.24% and 0.08% for Pincourt respectively (Table 2).

### 3.5. Contribution to substrate consumption

Beyond calibrating the immigration efficiency and the effective OHO population size in the influent, the analytical framework presented herein allows an initial estimation of the contribution to carbonaceous substrate conversions by low or negative net growth rate genera. The relative proportion of substrate consumed by each genus is given by Eq. 11 and Eq. 12. When the immigration efficiency is 1, the mass of the genus in the activated sludge is only contributed by the influent microbes; it does not increase due to substrate consumption, but only decreases due to decay. Genera with negative net growth rates (below the line of *μ*_*OHO,Net,i*_ = 0 in Figure 5) were considered to have zero contribution to the consumption of substrates. However, if *m*_*i*_ < 1 (below *μ*_*OHO,Net,i*_ = 0 and above *m*_*i*_ = 1), these genera could grow slowly and contribute to substrate consumption; although, the extent of this contribution remains to be evaluated.

The accumulation curves of resource consumption by genera ranging from highest (1) to lowest (0) immigration are shown in Figure 6, along with the accumulated relative abundances. Using the substrate consumption model (Eq. 12) and the data obtained in the current study, it can be estimated that 2.4% and 2.9% of the carbonaceous substrate resources were consumed by heterotrophic genera with zero or negative net growth rates in LaPrairie and Pincourt respectively (Figure 6a-b). If using the lysis-growth ASM1 model, the substrate consumption percentages are 4.5% and 5.4% in LaPrairie and Pincourt respectively (Figure 6c-d). These data suggest that genera subjected to significant immigration efficiencies still have important contributions to the substrate consumption in conventional activated sludge systems. Although, these data should be carefully evaluated given that the yield and decay rates can vary between taxa.^39^ In systems that heavily impacted by immigration, the relative abundance and substrate consumption of the slow-growth taxa can contribute largely to the microbial community assembly and functionality. The model framework should not only apply to heterotrophs described in this study, but also to autotrophs with proper assumptions made on growth and nutrient utilisation. On the other hand, heterotrophic populations can be differentiated into distinct groups co-existing in one system^40-43^ and the model can be further refined.

**Figure 6.**
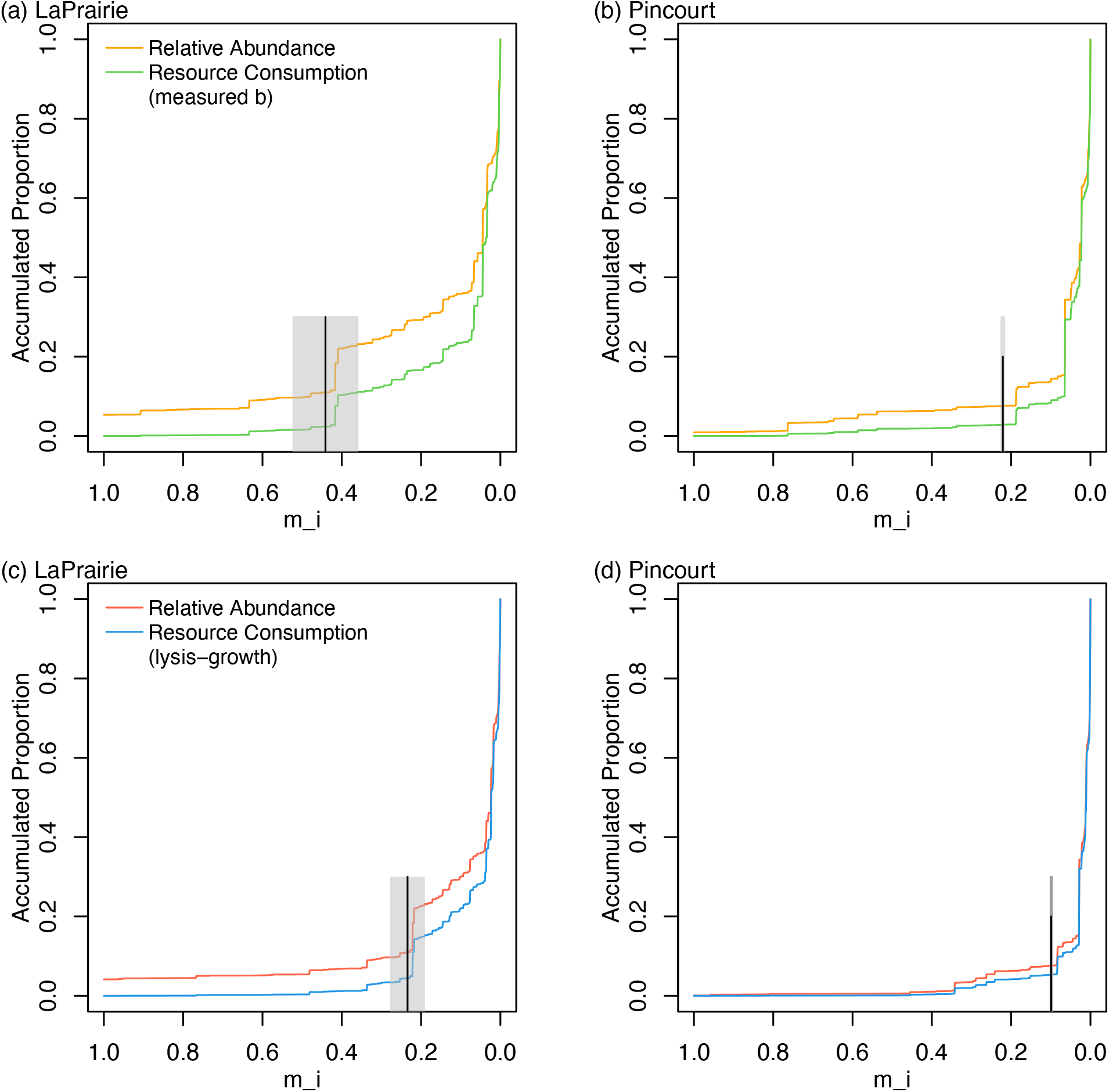
Accumulated relative abundance and proportion of substrate consumed by genera of different immigration efficiency in LaPrairie (a) and Pincourt (b) using influent and activated sludge microbial communities. Vertical lines indicate zero net growth rate reference lines. Shaded areas indicate 95% of confidence intervals.

## 4. Conclusions

Proper quantification of influent heterotrophic organisms is necessary for detailed and precise wastewater activated sludge modeling practice. Influent total biomass fraction determined using DNA mass is reliable and faster compared with conventional respirometry tests. Influent microbial community showed growth of certain genera in respirometry tests, such as *Acinetobacter, Rheinheimera*, and *Aquabacterium*. Compared to the conventional lumped heterotrophic population modelling in ASMs, specific taxa modelling using relative abundances based on 16S rRNA gene sequencing provides detailed information of taxon-specific mass-flow immigration efficiencies, growth rates and carbonaceous substrate consumption fractions. The mass-flow immigration models developed in this study showed that microbial genera varied in immigration efficiencies and growth rates. The genera with zero or negative net growth rates could not be neglected for their role on influent COD consumption (2.4% - 5.4% of total COD in two AS study sites). The detailed growth dynamics of influent heterotrophs could be incorporated in precise modeling of activated sludge with future efforts. The mass-flow immigration model has wide application to various microbial communities.

## Acknowledgement

The authors would like to thank the LaPrairie and Pincourt wastewater treatment plant operators for granting access to their wastewater facilities and helping with sampling. This work was supported by Discovery Grants from the Natural Science and Engineering Research Council of Canada RGPIN 341393-11 and RGPIN-2016-06498.

