## Supplementary for "Wastewater Influent Microbial Immigration and Contribution to Resource Consumption in Activated Sludge Using Taxon-Specific Mass-Flow Immigration Model"

**This document contains 8 pages, including texts for materials and methods, 4 supplementary figures, and 3 supplementary tables.**

### Materials and methods

#### Decay rate test

Decay rates were determined by respirometry test using a model AER-200 respirometer (Challenge Technology, USA) at 20 °C for 96 hours with triplicates. The working volume of each reactor is 800 mL with aerated activated sludge mixed liquor from each plant. Allylthiourea was added at 5 mg/mL as nitrification inhibitor. Eq. S1[29] was used to estimate $b_H$ and $X_{B,H,TO}$.

$$OUR|_t = (1 - f_D)b_H \cdot X_{B,H,TO} \cdot i_{O/XB,T} \cdot e^{-b_H t} \quad \text{(Eq. S1)}$$

Where OUR is the oxygen uptake rate, $f_D$ is the fraction of active biomass contributing to biomass debris assumed as 0.2, $b_H$ is the decay rate for heterotrophs, $X_{B,H,TO}$ is the initial biomass concentration, $i_{O/XB,T}$ is the mass of COD per mass of biomass as 1.20g-COD/g-TSS.

The decay rate was 0.304 ± 0.002 $d^{-1}$ for Pincourt activated sludge and 0.234 ± 0.027 $d^{-1}$ for LaPrairie activated sludge. The active biomass fraction was 32.8 ± 0.2 % and 24.3 ± 0.4 % respectively. Rate estimation was corrected for 11 °C using $r = r_{11^{\circ}C} \times 1.029^{(T-11^{\circ}C)}$.

#### Decay rate in lysis-growth model ASM1

The lysis-regrowth approach in ASM1 uses a different decay rate ($b_L$) with the traditional approach ($b$ measured), which can be calculated using the following equation [29] :

$$b_L = \frac{b}{1-Y(1-f_D')} \quad \text{(Eq. S2)}$$

Same notations as in Eq. S1. $f_D'$ is the fraction of active biomass contributing to biomass debris assumed as 0.08.

#### Error propagation

For the $m_i = 1$ and $\mu_{OHO,Net,i} = 0$ lines (Eq. 8a and Eq. 10a), we use $\log\left(\frac{\theta_x}{\theta} \cdot \frac{f_{OHO,Capt}}{1+b_{OHO,i}\theta_x} \cdot \frac{X_{Tot,Inf} \cdot \gamma_{DNA,Inf}}{X_{Tot,ML} \cdot \gamma_{DNA,ML}}\right)$ and $\log\left(\frac{\theta_x}{\theta} \cdot \frac{f_{OHO,Capt} X_{Tot,Inf} \cdot \gamma_{DNA,Inf}}{X_{Tot,ML} \cdot \gamma_{DNA,ML}}\right)$, i.e. the intercept of the two lines respectively.

Assuming independence between variables, the standard deviation of a function *f* with variables *x*, *y*... can be calculated using:

$$SD_f = \sqrt{(\frac{\partial f}{\partial x})^2 \cdot SD_x^2 + \left(\frac{\partial f}{\partial x}\right)^2 \cdot SD_x^2 + \cdots}$$

Results:

| SD | $\log(\frac{\theta_x}{\theta} \cdot \frac{f_{OHO,Capt}}{1 + b_{OHO,i}\theta_x} \cdot \frac{X_{Tot,Inf} \cdot \gamma_{DNA,Inf}}{X_{Tot,ML} \cdot \gamma_{DNA,ML}})$ | | $\log(\frac{\theta_x}{\theta} \cdot \frac{f_{OHO,Capt} X_{Tot,Inf} \cdot \gamma_{DNA,Inf}}{X_{Tot,ML} \cdot \gamma_{DNA,ML}})$ |
|---|---|---|---|
| | ASM3 | ASM1 | same for ASM1-3 |
| LaPrairie | 0.063 | 0.068 | 0.015 |
| Pincourt | 0.005 | 0.058 | 0.005 |

For substrate consumption, the vertical line indicates zero net growth rate $\mu_{OHO,Net,i} = 0$, which gives the $m_i$ value in Eq. 9, standard deviation is propagated from standard deviation of decay $b$.

Table S1. Average characteristics of WRRFs operation during the sampling month.

| Plant Operation Data | **LaPrairie** | **Pincourt** |
|---|---|---|
| Sampling dates | 2018-04-04<br>2018-04-09 | 2018-03-28<br>2018-03-30 |
| Process | PFR | PFR |
| Proportion of COD by origin (Residential : Industrial) | 45%:55% | 90%:10% |
| SRT (d) | 7 | 15 |
| HRT (hr) | 15 | 8 |
| Flow ($m^3$/d) | 72,598 | 11,025 |
| Influent | | |
| COD (mg/L) | 406 | 135 |
| $BOD_5$ (mg/L) | 99 | 41 |
| TSS (mg/L) | 194 | 69 |
| $NH_4^+$ (mgN/L) | 9.6 | N/A |
| P total (mgP/L) | 4.1 | N/A |
| Effluent | | |
| COD (mg/L) | 18 | 20 |
| $BOD_5$ (mg/L) | 4 | 6.1 |
| TSS (mg/L) | 6.4 | 9 |
| $NH_4^+$ (mgN/L) | 0.9 | 11 |
| P total (mgP/L) | 1.5 | N/A |
| Aeration Tank | | |
| Dissolved Oxygen (mg/L) | 4.33 | 2.3 |
| MLSS (mg/L) | 2,056 | 2,543 |
| MLVSS (mg/L) | 1,583 | 2,270 |
| Temperature (°C) | 11 | 11 |

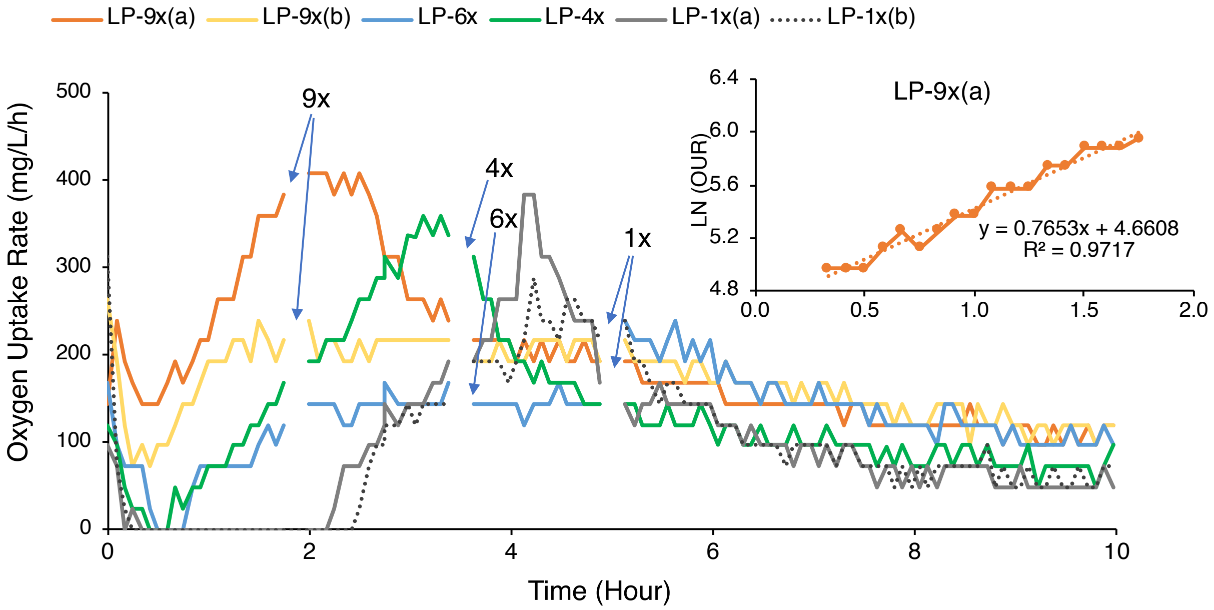


Figure S1. Oxygen uptake rate (OUR) curve of concentrated influent solids from LaPrairie. Sampling points of the biomass for determining the community composition are shown by arrows. Insert: The active biomass was calculated using the exponential growth phase.

Table S2. PCR primers and conditions for Illumina sequencing.

| | **Primer Pair with adaptor (5'-3')**[a] | **Conditions**[b] | **Cycle No.** |
|---|---|---|---|
| Illumina PCR1 | 515F (47 nt)<br>CTT TCC CTA CAC GAC GCT CTT CCG ATC TGT GYC AGC MGC CGC GGT AA<br>806R (54 nt)<br>GTG ACT GGA GTT CAG ACG TGT GCT CTT CCG ATC TGG ACT ACN VGG GTW TCT AAT | Initial denaturation: 94 °C, 3m<br>Denaturation: 94 °C, 30s<br>Annealing: 62 °C, 45s<br>Elongation: 72 °C, 1m<br>Final elongation: 72 °C, 10m | 25 |
| Barcode PCR2 | Uniprimer1 (45 nt)<br>AAT GAT ACG GCG ACC ACC GAG ATC TAC ACT CTT TCC CTA CAC GAC<br>Uniprimer2 (45 nt)<br>CAA GCA GAA GAC GGC ATA CGA GAT-variable index (8 nt)-GTG ACT GGA GTT C | Initial denaturation: 94 °C, 3m<br>Denaturation: 94 °C, 30s<br>Annealing: 59 °C, 20s<br>Elongation: 72 °C, 45s<br>Final elongation: 72 °C, 5m | 15 |

[a]Primers from IDT, Coralville, IA, USA
[b]Reagents from New England Biolabs Ltd, Whitby, ON, Canada: Taq DNA Polymerase with Standard Taq Buffer, Deoxynucleotide (dNTP) Solution Mix.

Table S3. Example of linear regression of Relative Abundance of Acinetobacter against growth factor in LaPrairie (LP) oxygen uptake rate (OUR) test at concentration factors of 1×, 4×, 6× and 9×, coded as dummy numbers 4, 3, 2, 1, as rank of F/M ratio respectively.

| Sample | Relative Abundance | Rank in Relative Abundance | Rank of F/M ratio |
|---|---|---|---|
| LP-OUR9x | 4.94% | 2 | 1 |
| | 4.28% | 1 | 1 |
| LP-OUR6x | 5.06% | 3 | 2 |
| LP-OUR4x | 11.92% | 4 | 3 |
| LP-OUR1x | 20.30% | 6 | 4 |
| | 12.94% | 5 | 4 |

Model Output

| | Coefficient | Std. Error | t value | Pr(>\|t\|) |
|---|---|---|---|---|
| Intercept | 0.2105 | 0.4658 | 0.452 | 0.67473 |
| Slope | 1.3158 | 0.1664 | 7.906 | **0.00138** |
| Adjusted $R^2$ | 0.92 | | | |

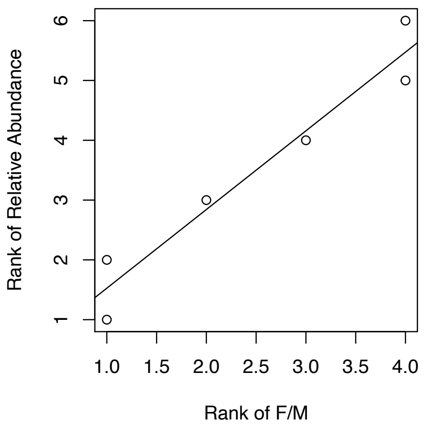


Figure S2. Example of linear regression of relative abundance of Acinetobacter against rank of F/M ratio in LaPrairie (LP) oxygen uptake rate (OUR) test at concentration factors of (1×, 4×, 6×, 9×).

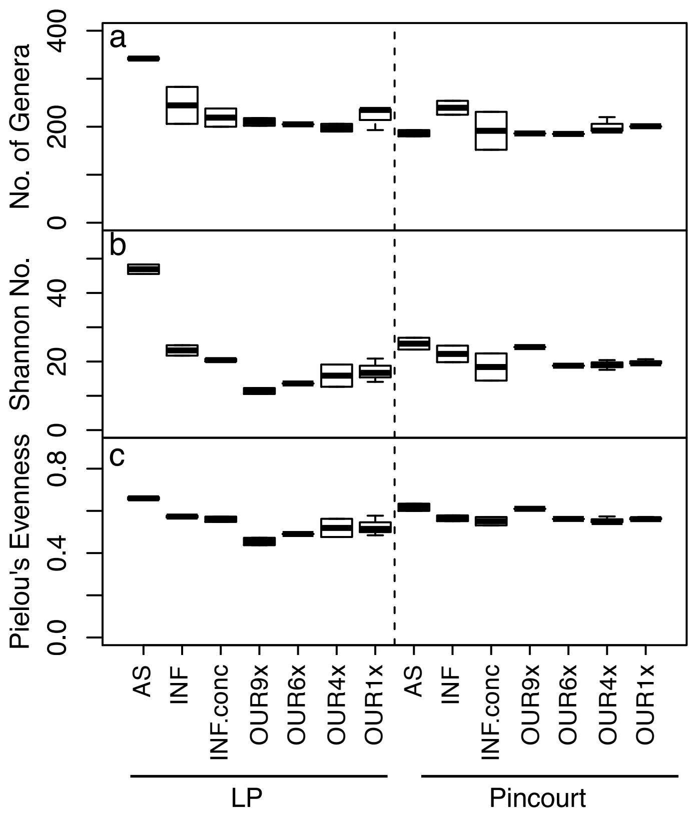


Figure S3. Alpha-diversity indexes at the genus level: (a) number of genera, (b) Shannon diversity number and (c) Pielou's evenness, in activated sludge (AS), influent (INF), concentrated influent solids (INF.conc), influent solids at the end of respirometry test (OUR) initiated with different solids concentration factors (1×, 4×, 6×, 9×).

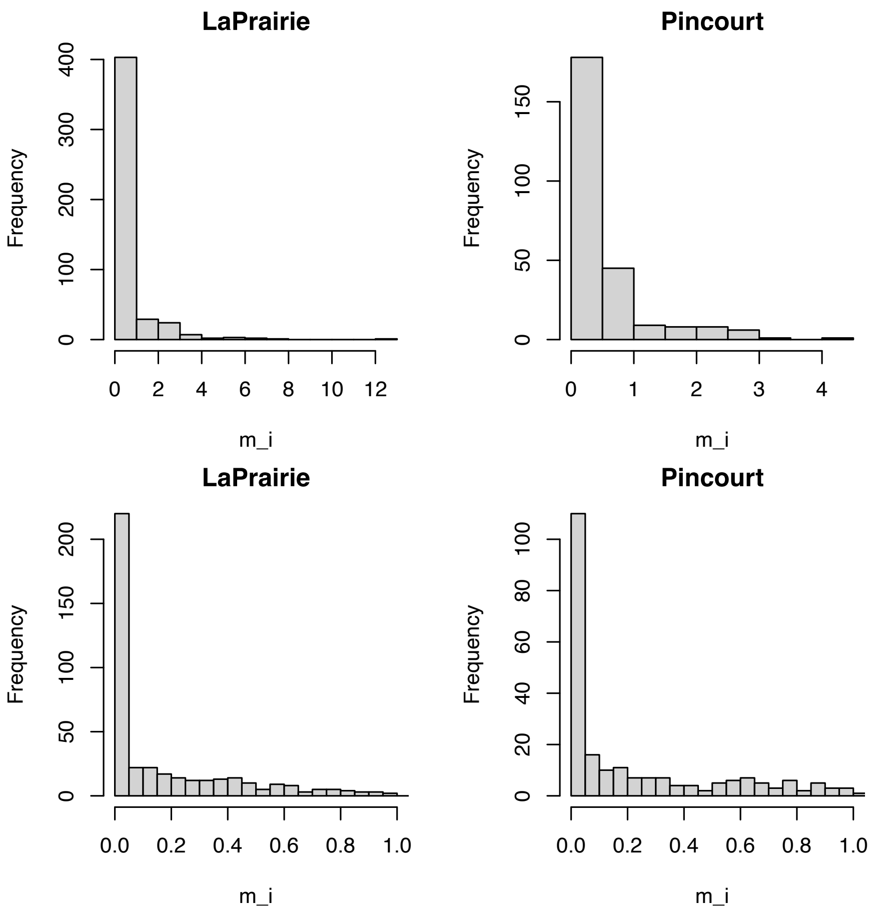


Figure S4. Histogram of immigration efficiencies in LaPrairie and Pincourt.
